# Transformation of plasmid DNA into *Prochlorococcus* via electroporation

**DOI:** 10.1101/2025.09.20.677525

**Authors:** Giovanna Capovilla, Kurt G. Castro, Christine A. Ziegler, Sallie W. Chisholm

## Abstract

*Prochlorococcus* is the most numerically abundant photosynthetic organism in the oceans and plays a role in global carbon cycling. Despite its ecological significance and the availability of over a thousand assembled genomes, progress in understanding gene function has been limited by the lack of genetic tools. Here, we report a reproducible electroporation-based protocol to introduce replicative plasmids into two strains of *Prochlorococcus* representing different ecotypes: MIT9313 (low-light adapted) and MED4 (high-light adapted). Using plasmids carrying a spectinomycin resistance cassette, we achieved transformation in ~33% of MED4 and ~10% of MIT9313 attempts, with greatest success when electroporating cells in late exponential phase. Transformed cells stably retained plasmids and expressed resistance genes, demonstrating functional uptake and gene expression. We also delivered a modified 13 kb plasmid carrying a CRISPR-Cpf1 system into MED4. While no targeted edits were observed, *cpf1* and *specR* were expressed, indicating successful delivery of large constructs and active transcription. These findings represent a key step toward genetic manipulation of *Prochlorococcus*, enabling future optimization of gene editing approaches and deeper functional analysis of its vast and largely uncharacterized pangenome.

## Introduction

*Prochlorococcus* is the smallest and most abundant photosynthetic organism in the oceans^1,2^. Despite the lack of genetic tools to manipulate this organism, metagenomic, metatranscriptomic, and metaproteomic studies of ocean samples have led to substantial progress in our understanding of *Prochlorococcus* biology and ecology, making it one of the best-described marine microorganisms^3–7^. *Prochlorococcus* has high photosynthetic efficiency^8,9^, an extremely small and compact genome^10,11^, is widely distributed, can account for 50% of the chlorophyll in oligotrophic ocean regions^8,9^, and is responsible for around 8.5% of global marine primary productivity^12^. *Prochlorococcus’* numerical predominance in oligotrophic oceans is due to its small size, which affords a heightened surface area-to-volume ratio, bolstering its access to essential nutrients^13^. Moreover, the vast genomic diversity observed within *Prochlorococcus* allows for the ecological niche dimensions of the collective *Prochlorococcus* population to expand beyond that of the individual lineage^9,11,14–20^. *Prochlorococcus* comprises a diversity of ecotypes characterized by distinct yet overlapping spatial distributions along environmental gradients in their natural habitat^1^. Each ecotype harbors a conserved “core genome” of roughly 1,000 genes, predominantly encoding essential housekeeping proteins. The remaining 700-1,400 “flexible” genes are heterogeneous among ecotypes, shaping the relative fitness of each ecotype along environmental gradients^1,21,22^. For each *Prochlorococcus* genome analyzed, either from cultured strains or wild single-cells, roughly 100 new genes are added to the *Prochlorococcus* pangenome — i.e. the complete set of genes found in all cells globally. Thus far, roughly 30,000 genes have been identified, most of which are of unknown function; the global *Prochlorococcus* pangenome is projected to consist of greater than 80,000 genes^1^.

Over a thousand complete or nearly complete genomes are available for analysis^1,19,23^, and culturable natural ecotypes exhibiting differences in the presence/absence of specific genes give us clues as to the selective pressures shaping the distributions of genes of known function, such as iron uptake^24^, phosphorus uptake^25,26^, nitrogen sources^27,28^, horizontal gene transfer^29^, mixotrophy^30^, and chitin utilization^31^, to cite a few, have been unraveled. However, our inability to assign functions to the numerous unannotated genes in these cells obscures many potentially illuminating niche dimensions. Furthermore, aside from its significant ecological and biogeochemical importance, the biology of *Prochlorococcus* is of interest as a potential chassis for solar-driven synthetic biology^32^. This microorganism represents nature’s simplest photosynthetic “machine,” making it an appealing organism for manipulation^32–35^.

Despite considerable interest, progress in developing a robust genetic system has been a challenge because of intrinsic obstacles, such as *Prochlorococcus*’ slow growth rate, their reluctance to grow axenically on solid media, the inefficient methods of recovering from single cells such as dilution-to-extinction techniques, specific media requirements^36^, and sensitivity to trace metal contamination^37,38^ and reactive oxygen species^39^. A successful *Prochlorococcus* genetic transformation has been reported based on a conjugation from an *E. coli* donor strain to a low-light adapted strain of *Prochlorococcus*, MIIT9313^40^ and a similar technique is routinely used to transform *Synechococcus* marinus, closely related to *Prochlococcus*^*41*^. However, this multi-step and relatively time-consuming technique, which necessitated at least six months for completion^40^, has not been successfully replicated in *Prochlorococcus*^*42*^ since, revealing it to be an inefficient and unreliable method for this genus.

In the past decade, we have put tremendous effort into building a genetic system for *Prochlococcus*, focusing on the use of electroporation as an alternative strategy for delivering DNA. Using fluorescein-labeled oligonucleotides as a probe for DNA delivery and a dead cell staining strategy, we quantified cells that survived electroporation and identified the optimal electric field and time of electroporation^42^. In the work reported here, we build upon these findings to successfully transform both a low- and high-light adapted *Prochlorococcus* strain with replicative plasmids that confer resistance to an antibiotic.

## Results and Discussion

### Streamlining of the protocol

Our published electroporation protocol enabled the identification of fluorescein-labeled oligonucleotides in living cells in a liquid medium via flow cytometry^42^. However, transformation of fluorescein-labeled plasmids could not be tested due to technical constraints, and no transformants were recovered in selective plates^42^. It is noteworthy, however, that growth of *Prochlorococcus* even on non-selective plates is notoriously challenging^36^. The cells typically require a heterotrophic ‘helper strain’ which produces catalase^43^, detoxifying reactive oxygen species and promoting *Prochlorococcus* growth by reducing oxidative stress^39^. Using this method of culturing *Prochlorococcus* on solid media for our purposes would therefore necessitate subsequent re-purification to obtain axenic *Prochlorococcus* mutants, representing a significant bottleneck.

Alternative strategies have been suggested, such as recovering axenic *Prochlorococcus* cultures in a semi-solid 0.3% agar solution supplemented with pyruvate, which helps to quench reactive oxygen species^42,44^. We have succeeded in using this approach for recovering and selecting *Synechococcus* transformant cells after electroporation^45^. However, while wildtype *Prochlorococcus* cells can grow on non-selective semi-solid agar plates (albeit with colonies typically only becoming visible to the naked eye after 2-3 months)^42^, we did not succeed in recovering electroporated cells with selective plates using this approach.

To avoid the complexities with plating, we opted to maintain all cultures in liquid media to assess the potential for *Prochlorococcus* cells to recover from electroporation in the work reported here. Our objective was to determine whether replicative plasmids could be introduced into *Prochlorococcus* cells and whether transformants could recover under selective pressure by expressing an antibiotic resistance cassette encoded within the plasmids. We identified spectinomycin resistance as a suitable selectable marker since kanamycin, routinely used for various marine *Synechococcus* strains, was ineffective in suppressing the emergence of spontaneous kanamycin-resistant colonies at high frequencies^42^.

Briefly, our transformation protocol is as follows. *Prochlorococcus* cultures grown in amended seawater medium were collected and washed twice with ice-cold mannitol to remove residual salts (Fig. 1a; see Methods for details) and concentrate cells. Plasmid DNA was then added to the concentrated cells, followed by electroporation. Cells were subsequently transferred to fresh media for a 24-hour recovery period prior to antibiotic selection (Fig. 1a).

**Figure 1.**
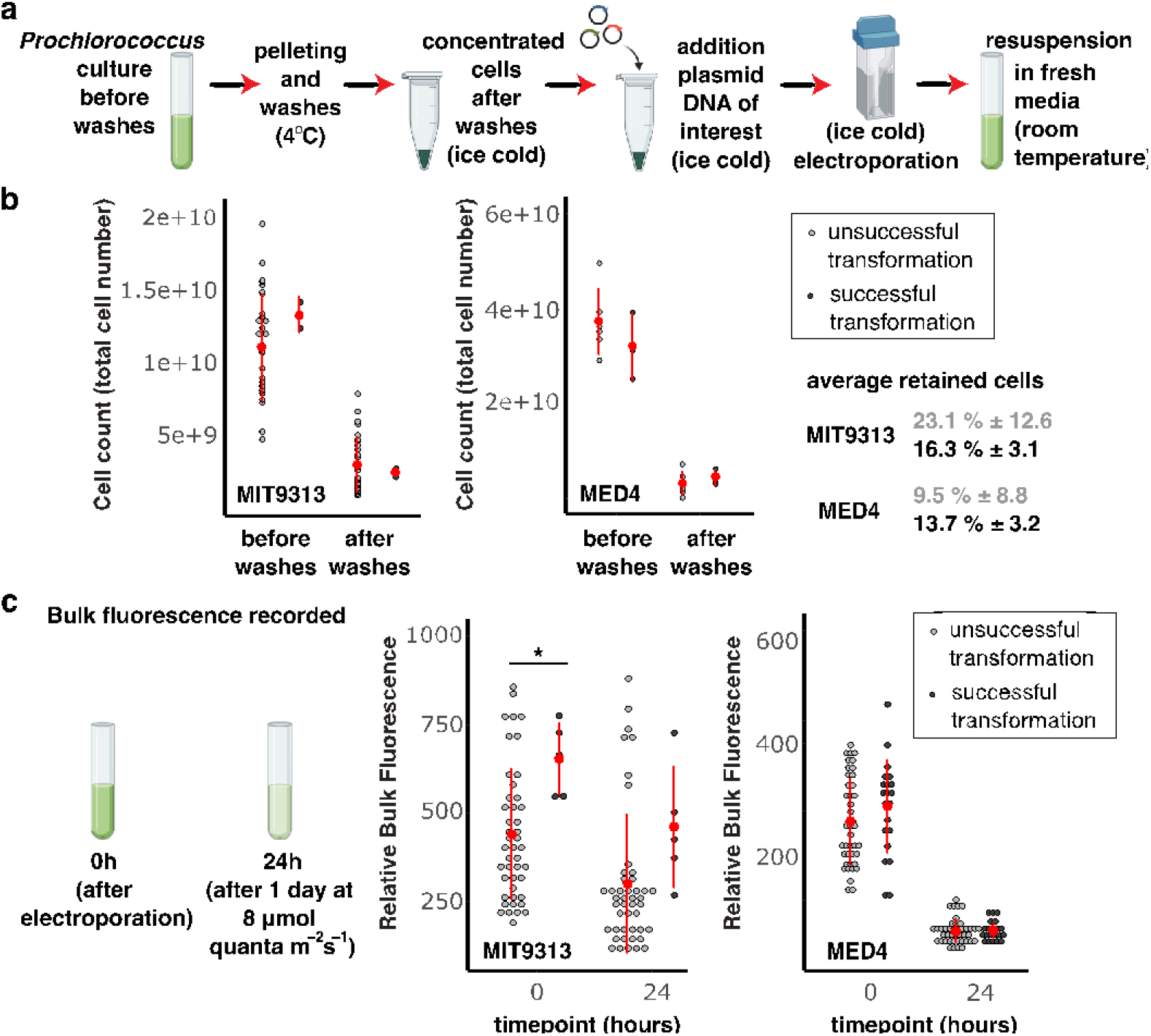
Overview of the electroporation protocol. **a**. Schematic representation of the steps in the protocol. *Prochlorococcus* cells are grown in seawater media, pelleted, and washed twice in osmoprotectant to remove salt traces without lysing the cells. DNA plasmids are added to the ice-cold concentrated cells. Samples are electroporated and resuspended in fresh seawater medium. **b**. Cell number was measured in a subset of samples before and after washes to determine the number of cells lost during the two washes, which is reported in percentages. Unsuccessful attempts are shown in grey, and data relative to successful transformants is shown in black. Average and standard deviation are shown in red. **c**. Bulk fluorescence was recorded immediately after electroporation and after 24 hours incubation (*p < 0.05 using Welch’s t test).

### Testing Protocol Success Rate

To investigate the protocol success rate in delivering DNA plasmids into *Prochlorococcus* cells we focused on two strains: low-light–adapted strain MIT9313 and high-light–adapted strain MED4. A total of 50 transformation attempts were performed for MIT9313 and 63 for MED4. Transformation was considered successful when cells that carried a plasmid conferring antibiotic resistance recovered under antibiotic selection. Of the attempts, 5/50 (10%) were successful for MIT9313 and 21/63 (~33%) for MED4. These experiments allowed us to evaluate protocol success rate across multiple parameters and identify the conditions necessary for effective transformation.

To assess cell loss during the washing steps, we monitored cell numbers in a subset of samples before and after the two washes (Fig. 1b). As previously reported, up to 90% of cells were lost during this process^42^. However, the extent of cell loss did not differ significantly between successful and unsuccessful transformation attempts.

We also measured bulk fluorescence of electroporated cultures immediately after transfer to fresh medium (t= 0h after electroporation) and again after 24 hours of recovery, prior to antibiotic selection. In all samples we observed a drop in fluorescence between the two time points (Fig. 1c). In MIT9313, successful samples displayed higher fluorescence immediately after electroporation compared to unsuccessful samples; however, this difference was not maintained after the recovery period and was not observed in MED4 (Fig. 1c). Therefore, changes in bulk fluorescence of the cultures do not appear to be a reliable indicator of transformation success.

After 24 hours of recovery from electroporation, cells were distributed into tubes for antibiotic selection (Fig. 2a). To monitor overall cell viability, 10% of the culture was reserved as an experimental control, with no antibiotic added, while the remaining 90% was subjected to selection by spectinomycin (Fig. 2a). Although cells were unevenly distributed between the control and selection conditions, fresh medium was added to equalize the final volume.

**Figure 2.**
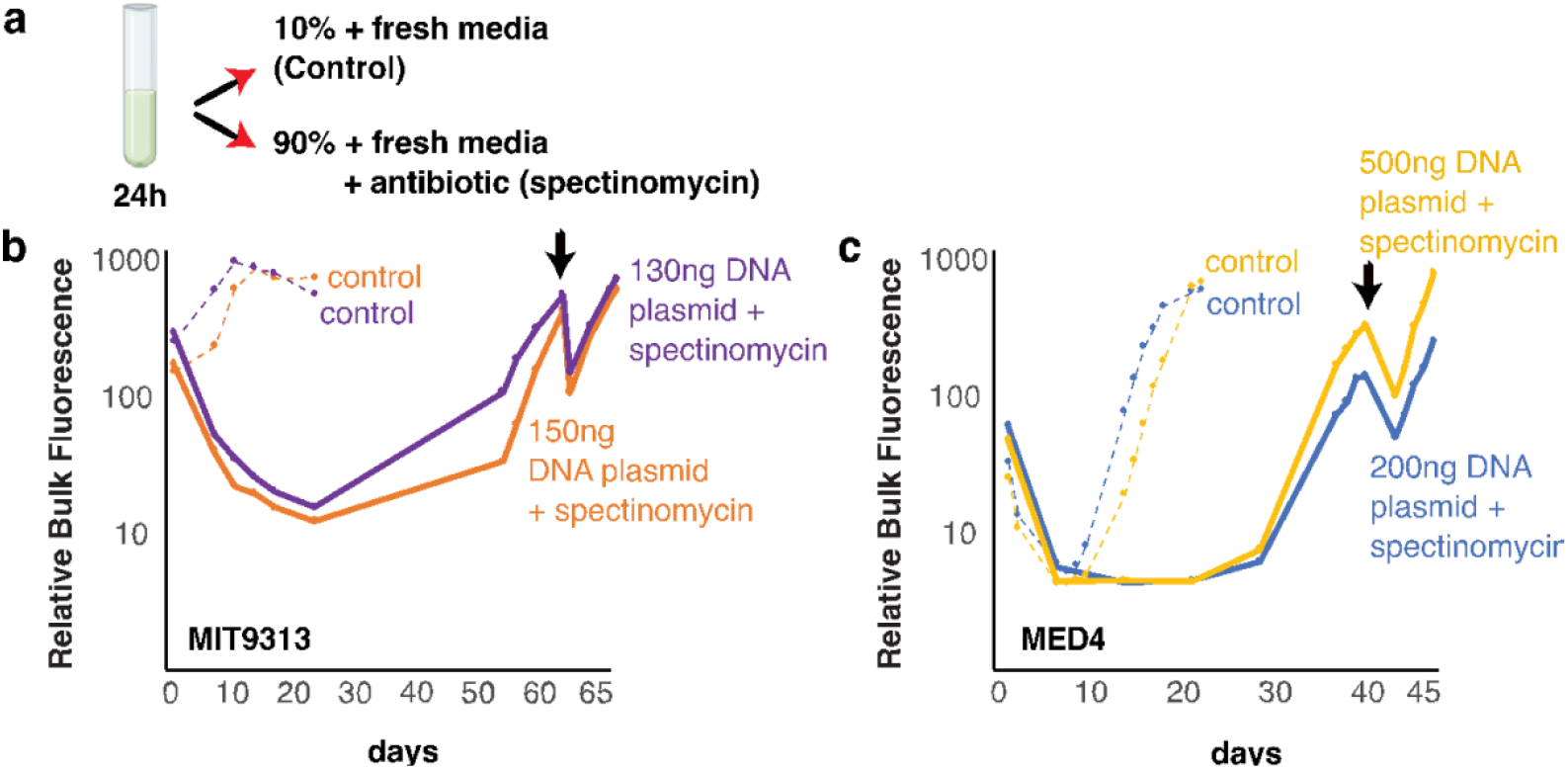
Recovery of successful transformants. **a**. Cells are unevenly distributed into two tubes, to which fresh medium is added to keep the final volume even. Spectinomycin is added to 90% of cells, while 10% of cells are kept without antibiotics as a control to ensure cells have survived electroporation and washes. Growth was monitored by bulk chlorophyll fluorescence. Two successful examples for each ecotype used in this work: **b**. MIT9313 (Low light adapted ecotype), and **c**. MED4 (High light adapted ecotype). Dashed lines show controls, and bold lines show growth of the transformants under antibiotic selection. The amount of DNA replicative plasmid with a spectinomycin resistance cassette electroporated in each sample is reported in brackets. Black arrows indicate the first transfer of transformants in fresh media with selection.

Control samples (no antibiotic) recovered rapidly and showed visible growth within 10 days (Fig. 2b–c), confirming that cells survived electroporation and washing steps. In contrast, samples under antibiotic selection exhibited a consistent drop in fluorescence, likely reflecting the death of most cells. Over time, a small subpopulation — those that successfully acquired the plasmid — began to recover and grow to detectable levels (Fig. 2b–c).

For the high-light–adapted ecotype MED4, the most reliable predictor of successful transformation was the recovery time of control samples (no antibiotic) following electroporation (Fig. 3a). Controls from successful transformations consistently recovered significantly faster than those from unsuccessful attempts. This trend was not observed in the low-light–adapted strain MIT9313, which may reflect fundamental physiological differences between the two ecotypes. However, it is also possible that the lack of a clear pattern in MIT9313 is influenced by the limited number of successful transformations in this strain.

**Figure 3.**
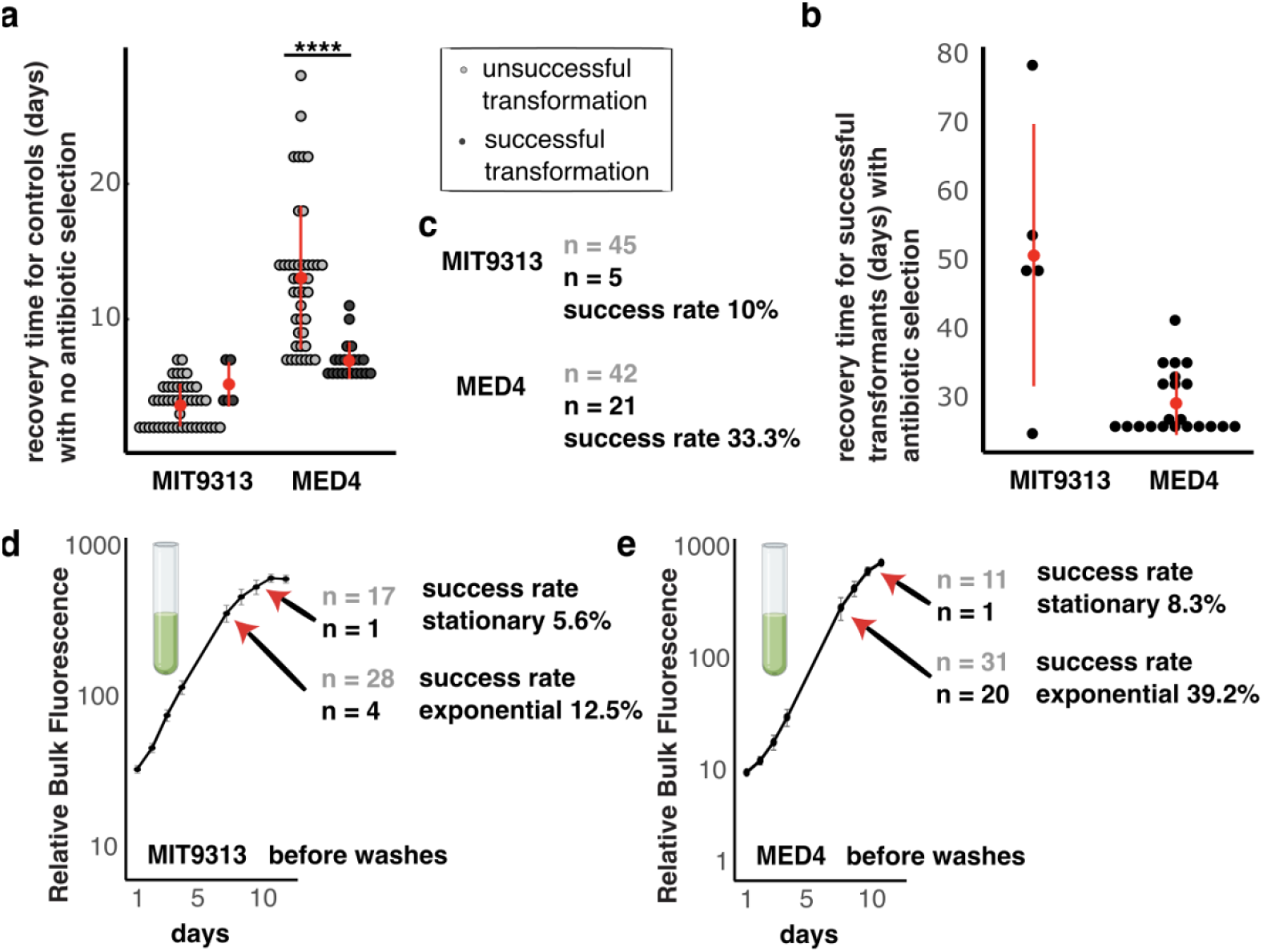
Transformation success rate. **a**. Bulk fluorescence was recorded regularly in both controls and samples under antibiotic selection. Recovery day of the controls, calculated as the first day fluorescence started increasing, is reported for both ecotypes. Unsuccessful attempts are shown in grey, and data relative to successful transformants is shown in black. Average and standard deviation are shown in red. (****p < 0.0001 using Welch’s *t* test). **b**. Recovery time of the successful transformants is reported for each ecotype. Average and standard deviation are shown in red. **c**. Success rate is reported in percentage. Bulk fluorescence of the cells used for transformation for MIT9313 and MED4 is shown in **d**. and **e**., respectively, and reported in percentage. Success rate is calculated separately for samples in which cells were harvested in exponential or early stationary phases. Unsuccessful attempts are shown in grey, and data relative to successful transformants is shown in black.

**Figure 4:**
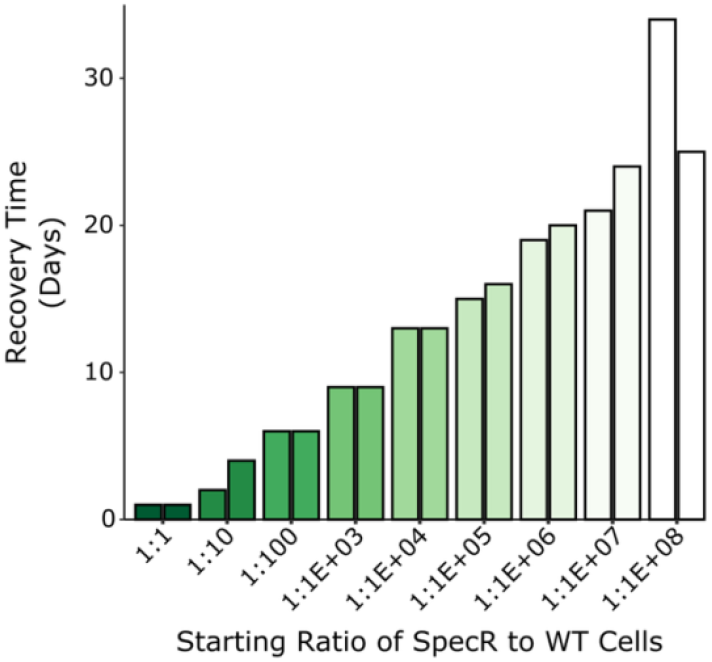
The ratio of transformed cells to wildtype cells impacts the rate of recovery. Keeping the total number of cells constant, various ratios of SpecR to WT cells were combined and subjected to spectinomycin. Recovery time shows a linear response to the ratio of transformed cells to WT cells.

### Protocol optimization

Overall, transformation was successful in 5 out of 50 attempts (10%) for MIT9313 and in 21 out of 63 attempts (~33%) for MED4. To optimize the protocol, we investigated whether the growth phase at the time of cell harvest influenced transformation success rate. Given the substantial cell loss during the seawater removal washes prior to electroporation, we focused on harvesting cells during the late exponential and early stationary phases to maximize initial cell numbers (Fig. 3d–e).

When analyzing success rates by growth phase, we observed that cells harvested in early stationary phase had significantly lower transformation success rates than those harvested in late exponential phase. For MIT9313, only 1 of 18 stationary-phase samples (5.6%) was successful, and for MED4, just 1 of 12 (8.3%) (Fig. 3d–e). In contrast, late exponential-phase cells showed a higher transformation efficiency: 4 of 32 samples (12.5%) were successful in MIT9313, and 20 of 51 (39.2%) in MED4 (Fig. 3d–e). Thus, harvesting cells in the late exponential phase improves transformation success rate.

### Calculating transformation efficiency

We next wondered if we could measure the transformation efficiency of *Prochlorococcus* electroporation in our successful experiments, and how this affects the rate of recovery. For this set of experiments, we combined cultures of wildtype (WT) MED4 with cultures of MED4 that had successfully been transformed with pJS1, a plasmid containing a spectinomycin-resistance (SpecR) marker, in various ratios. Testing 10-fold dilutions between 1:1 and 1:1×10^8^ SpecR:WT cells, we noticed a clear linear relationship between the recovery time and fraction of cells containing the antibiotic resistance plasmid. While electroporation itself may cause stress to *Prochlorococcus* cells, cultures can still recover successfully, even in situations where just 3.5 cells in a pool of 3.5×10^8^ cells/mL contain the antibiotic resistance plasmid. Furthermore, this recovery time is consistent with what we see for electroporation and recovery of MED4 cultures, indicating the transformation efficiency for *Prochlorococcus* MED4 electroporation is around 1×10^−8^. Note that we see the largest variability in the recovery time in the 1:1×10^8^ condition because so few cells (3 or 4) contain the SpecR marker in the inoculum and is therefore subject to more stochasticity in starting conditions.

### Testing plasmid uptake and gene expression in transformed *Prochlorococcus* cells

A variety of plasmids carrying the same antibiotic resistance cassette were used across electroporation attempts, with varying amounts of DNA per sample (Dataset S1, Table S1). The smallest plasmid used was *pJS1*, a 6kb autonomously replicating empty vector carrying a *SpecR*^*46*^. To confirm plasmid uptake, we performed PCR on both concentrated, lysed cells and the surrounding cell-free medium. As expected, amplification products were detected from lysed cells, confirming that the plasmids were successfully introduced (Fig. 5a). Interestingly, faint amplification signals were also observed from the cell-free medium, suggesting that some cells may have lost the plasmid during growth—potentially through cell lysis and plasmid release into the medium. This assay was performed after several passages in fresh medium, suggesting that any remaining signal is unlikely to originate from residual plasmid introduced during electroporation, but rather from plasmid released by transformant cells.

**Figure 5.**
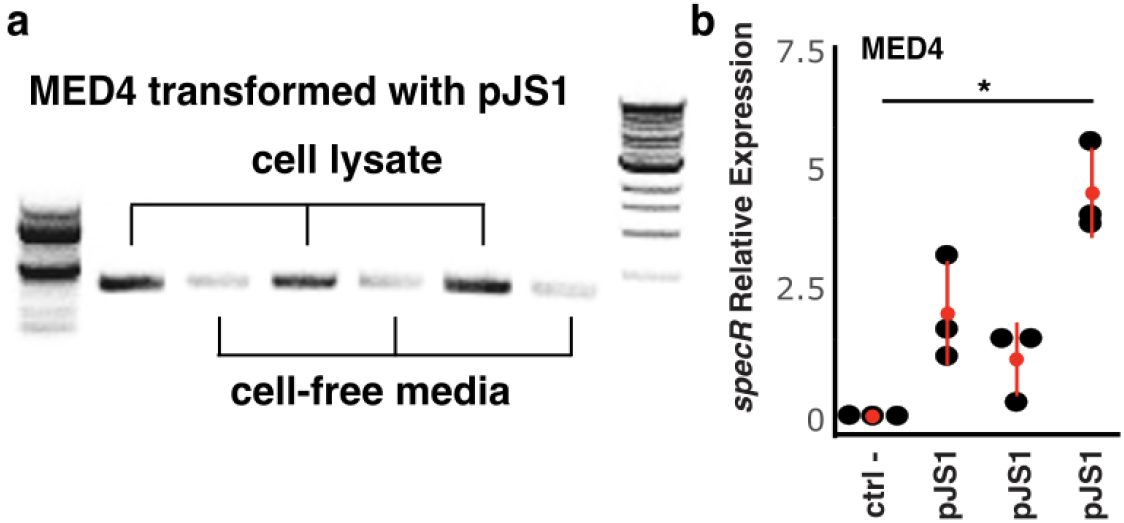
DNA plasmids transformed into *Prochlorococcus* cells. a. PCR products indicate amplification of DNA plasmid pJS1 in 3 independent replicates. Cell lysates and cell-free media have been tested for each. 100bp and 1kb markers are shown on the left and right sides. **b**. Expression (measured by qPCR) of the spectinomycin resistance cassette in wild-type (negative control) and mutant lines in mid-exponential growth in relation to the housekeeping gene, *rnpB*, in natural seawater-based Pro99. Average and standard deviation are shown in red. (*\*P < 0*.*05* using Welch’s *t* test). Primers used for the amplifications are listed in Table S2.

To further confirm that transformed cells could utilize the genetic information carried by the plasmids, we assessed expression of the spectinomycin resistance gene (*specR*) using quantitative PCR, with *rnpB* as a housekeeping control. Expression of *specR* was detected in all samples transformed with plasmid *pJS1* (Fig. 5b), demonstrating that cells not only received the plasmid but were also capable of transcribing the resistance gene.

We also tested several fluorescent proteins—GFP, mCherry, and the orange fluorescent protein CyOFP1—with the goal of generating fluorescently labeled *Prochlorococcus* cells (Dataset S1, Table S1). Successful expression would enable real-time monitoring of co-cultures involving different *Prochlorococcus* strains. Although some plasmids carrying these fluorescent genes were successfully introduced into cells– as confirmed by spectinomycin resistance and recovery – no fluorescent signal above background autofluorescence was detected via flow cytometry. To enhance expression, we replaced the T7 promoter with the strong endogenous *Prochlorococcus* promoter *psaA*. However, even in these constructs, no detectable fluorescence was observed in any of the successful transformations. These results suggest that, despite plasmid uptake, use of fluorescent proteins in *Prochlorococcus* remains challenging—potentially due to improper protein folding, poor expression, low signal, or detection interference due to cellular pigments.

In our previous work in *Synechococcus*^*45*^, we successfully delivered the ~13 kb plasmid *pSL2680*—carrying a *CRISPR-Cpf1* cassette—into *Synechococcus* WH7803 via electroporation. That plasmid, which includes a kanamycin resistance cassette, was effectively used to generate loss-of-function mutants^45^. To test whether *Prochlorococcus* MED4 could similarly take up and utilize a larger plasmid, we modified *pSL2680* by replacing the kanamycin resistance cassette with the same spectinomycin resistance cassette in *pJS1*. The modified *pSL2680-Spec* plasmid was successfully electroporated into MED4 cells, and we used it to attempt CRISPR-based gene editing.

We designed seven constructs (Dataset S1, Table S1) targeting five non-essential genes, aiming to induce deletions or point mutations without lethal effects. These constructs included guide RNAs (sgRNAs) targeting specific genes, and in some cases, a homologous repair template to enable precise genome editing via homologous recombination. Of the seven constructs, three were successfully delivered into cells: one targeting glucose-inhibited division protein A, and two targeting molybdenum cofactor biosynthesis protein— one with a repair template and one without.

Expression of the *cpf1* nuclease gene was confirmed in all three transformed samples (Fig. S1). In one sample, *cpf1* expression was notably low, potentially due to spontaneous mutations, as *specR* expression was also reduced (Fig. S1a). The other two samples showed robust expression of *cpf1* (Fig. S1b). However, Sanger sequencing of the transformed samples DNA did not reveal any mutations or deletions in the target genes. It is possible that the introduction of double-strand breaks in the *Prochlorococcus* genome at these loci is lethal and was counter-selected by spontaneous mutations.

## Conclusions

This work represents a significant step forward in developing a genetic system for *Prochlorococcus*, a long-standing goal in the field. Our results highlight key differences in how two diverse *Prochlorococcus* strains respond to washing and recovery steps— differences that may be due to their distinct physiologies. While MIT9313 cells were transformable via conjugation in 2006^40^, that protocol proved to be unreproducible in subsequent years. The results presented here show potentially relevant low transformation success rates for MIT9313, while a much more reliable performance was seen with MED4.

To further improve this protocol, additional strategies should be explored. For example, chitin has been used to induce natural competence in *Vibrio* spp., and our transcriptomics analyses showed that genes involved in horizontal gene transfer are upregulated in low light *Prochlorococcus* in the presence of chitosan^31^. Such findings suggest that chitin derivatives may enhance transformation efficiency and success rate.

In this study, we also demonstrate that *Prochlorococcus* MED4 can take up large plasmids carrying CRISPR-Cpf1 constructs and express the nuclease. Although no gene edits were detected, this shows that core components of the CRISPR system can be introduced and expressed. The lack of detectable deletions may be due to suboptimal guide RNA design, improper folding or activity of the Cpf1 protein, or the generation of lethal double-strand breaks. Further work is needed to optimize guide selection, assess nuclease function, and validate CRISPR as a genome-editing tool in *Prochlorococcus*. However, given its success in *Synechococcus*^*45*^, there is good reason to believe this approach can be refined.

With plasmid delivery now achievable, other genetic strategies—such as transposon mutagenesis—can also be considered once transformation efficiency is improved. Additionally, implementing methods to isolate clonal populations, such as filter plating^47^ or dilution-to-extinction protocols, could greatly improve downstream selection and analysis, particularly given the challenges of working with *Prochlorococcus* on traditional agar plates.

## Materials and Methods

### Culture conditions and growth curves

*Prochlorococcus* cells were grown under constant light flux at 12 μmol quanta m^−2^ s^−1^ (MIT9313) or 30 μmol quanta m^−2^ s^−1^ (MED4) at 24°C in natural seawater-based Pro99 medium containing 0.2-μm-filtered Sargasso Sea water, amended with Pro99 nutrients (N, P, and trace metals) prepared as previously described36.

Growth was monitored using bulk chlorophyll fluorescence measured with a 10AU fluorometer (Turner Designs). Cell concentration was measured using a Guava easyCyte 12HT flow cytometer (EMD Millipore, Billerica, MA). Cells were excited with a 488 nm laser for measuring chlorophyll fluorescence (692/40 nm).

### Electrocompetent cell preparation

For each sample, 50 mL of *Prochlorococcus* culture in late exponential or early stationary phase was harvested by centrifugation (10 min, 4°C, 8000 rpm) to remove seawater-based growth medium. The resulting cell pellet was gently resuspended in 10 mL of ice-cold 0.4 M mannitol (#63560 Merck Life Science UK Limited, Gillingham, Dorset, UK) supplemented with 1 mM HEPES (pH 7.5) (#H8651 Merck Life Science UK Limited, Gillingham, Dorset, UK), taking care to avoid drying the pellet. This washing step was repeated twice, with each wash followed by centrifugation (10 min, 8000 rpm, 4°C) and complete removal of the supernatant to eliminate residual salts.

Cell loss during washing was substantial, with up to 90% of the culture lost. MIT9313 cells exhibited a strong tendency to adhere to plastic surfaces, while MED4 formed loose pellets, resulting in significant loss during supernatant removal.

After the second wash, the final pellet was gently resuspended in 80 µL of ice-cold 0.4 M mannitol with 1 mM HEPES (pH 7.5). These concentrated cells were then immediately used for electroporation and kept ice-cold.

### Electroporation

Electrocompetent cells were prepared as described and kept on ice. Plasmid DNA (100 ng to 1 µg; see Dataset S1) was added to 80 µL of ice-cold cells, which were then transferred to pre-chilled 2 mm gap, sterile, electroporation cuvettes. Electroporation was performed using the following settings: 2.5 kV, 500 Ω, and 25 µF.

Immediately after electroporation, cells were gently resuspended in 1 mL of fresh, room temperature Pro99 medium, using the same medium batch in which cells were previously grown to minimize stress due to batch variability. The cell suspension was then transferred to 24 mL of fresh Pro99 medium for a total volume of 25 mL. Bulk fluorescence was recorded at this point. Tubes were incubated for 24 h under the same light intensity used for the pre-electroporation cultures. After incubation, bulk fluorescence was measured again. In all samples, a notable decrease in fluorescence was observed, in contrast to previous observations in *Synechococcus* WH7803 cultures^45^ following electroporation.

### Mutant selection

After 24 h of recovery without selection, cultures were split into a control (1/10 of the original volume) and a selection tube. Both tubes were brought to a final volume of 25 mL using fresh Pro99 medium. Spectinomycin was added to the selection cultures at a final concentration of 25 µg mL^−1^. Cell growth was monitored over time by measuring bulk culture fluorescence in both control and antibiotic-treated samples.

### Quantitative PCR analysis

*Prochlococcus* cells were collected by centrifugation. RNA samples were extracted with a standard acidic Phenol:Chloroform protocol and measured with Nanodrop (Thermo Scientific). RevertAid First Strand cDNA Synthesis Kit (Thermo Scientific) with random primers was used to obtain cDNA. Quantitative PCR reactions were performed in a CFX96 thermocycler (Bio-Rad) using the primers listed in Table S2. The expression of rnpB gene was used to normalize the results.

### Constructs design

To replace the resistance cassette in pSL2680, the plasmid was digested with *MluI-HF*® (NEB #R3198), and the digested fragment was purified using the NucleoSpin Gel and PCR Clean-up (Machery-Nagal #740609.50). The spectinomycin resistance cassette was PCR-amplified from plasmid pJS1, and the plasmid was reassembled using the NEBuilder® HiFi DNA Assembly Cloning Kit (NEB #E5520S). Assembly products were transformed into E. coli NEB® 5-alpha Competent Cells (NEB #C2987) following standard cloning procedures.

Similarly, to swap fluorescent protein genes, plasmid pJUMP^48^ was digested with *NotI-HF®* (NEB #R3189), and the GFP gene was replaced with either mCherry or CyOFP1, both amplified from plasmids provided by collaborators. Constructs were tested for fluorescence in *E. coli* using a Guava easyCyte 12HT flow cytometer (EMD Millipore, Billerica, MA) to confirm expression prior to electroporation into *Prochlorococcus*.

CRISPR vectors were built using the plasmid pSL2680 carrying a spectinomycin resistance cassette. Single guide RNAs (sgRNAs) were designed as previously described^45,49^. For constructs requiring homologous recombination, repair templates were synthesized as left and right homology arms (700–750 bp each) corresponding to the upstream and downstream regions of the target genes. Primer sequences used are listed in Table S2.

### Transformation Efficiency and Recovery Experiments

We inoculated 25mL of axenic MED4 *Prochlorococcus* cultures with a total of 3.5×10^8^ cells/mL in standard Pro99 media, as measured by flow cytometry. The ratio of cells containing the SpecR pJS1 plasmid (SpecR) to cells without the plasmid (WT) was altered in each condition such that we had 1:1, 1:10, 1:100, and continuing 10-fold dilutions up to 1:1×10^8^, which corresponded to 3.5 SpecR cells mixed with 3.5×10^8^ WT cells, and all cultures were subjected to 25 µg mL^−1^ spectinomycin. Cultures were monitored every other day for bulk relative fluorescence and were considered “recovered” when they hit mid-log phase in their growth curves (>200 relative fluorescence units) (Fig. S2). Each dilution was performed in biological duplicates.

## Acknowledgements

This research was supported in part by grants from the National Science Foundation (NSF-EDGE IOS-1645061 to S.W.C., NSF-EDGE IOS-2035181 to S.W.C. and Burton, B, and NSF-EDGE IOS-2319332 to S.W.C.) and from the Simons Foundation (Life Sciences Award IDs 337262, 647135, 736564 to S.W.C.; SCOPE Award ID 329108 and 721246 to S.W.C.). G.C. was supported by the Human Frontier Science Program (LT000069/2019-L). and C.A.Z. was supported by a Simons Foundation Postdoctoral Fellowship in Marine Microbial Ecology (LS-FMME-00003951). This paper is a contribution from the Simons Collaboration on Ocean Processes and Ecology (SCOPE).

## Supplementary Figures

**Figure S1:**
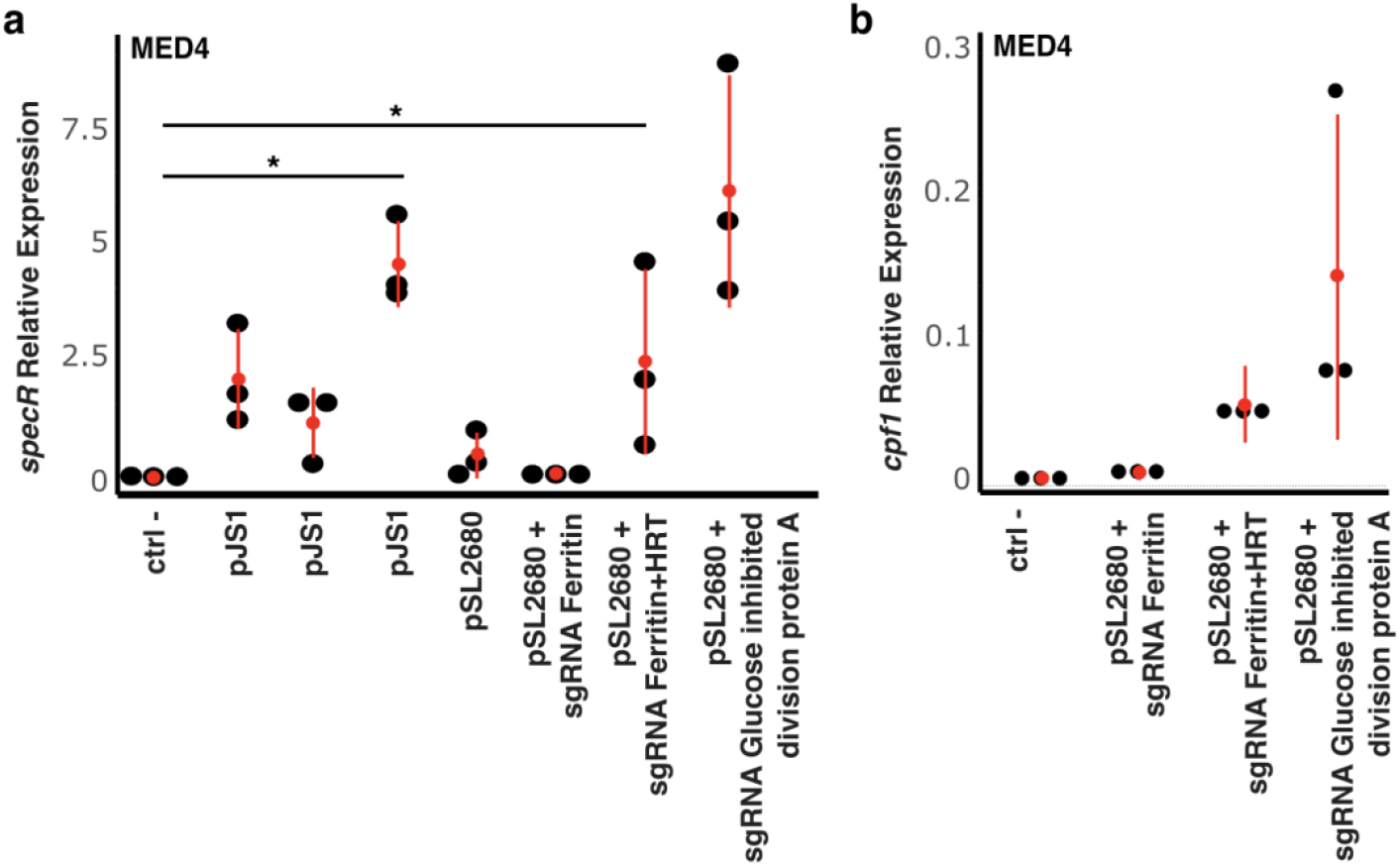
Expression, (measured by qPCR) of the spectinomycin resistance cassette **(a)** and Cpf1 **(b)** in wild-type (negative control) and mutant lines in mid-exponential growth in relation to the housekeeping gene, *rnpB*, in natural seawater-based Pro99 medium in absence (control) or presence of antibiotic selection (mutants). Average and standard deviation are shown in red. (*\*P < 0*.*05* using Welch’s *t* test). Primers used for the amplifications are listed in Table S2.

**Figure S2:**
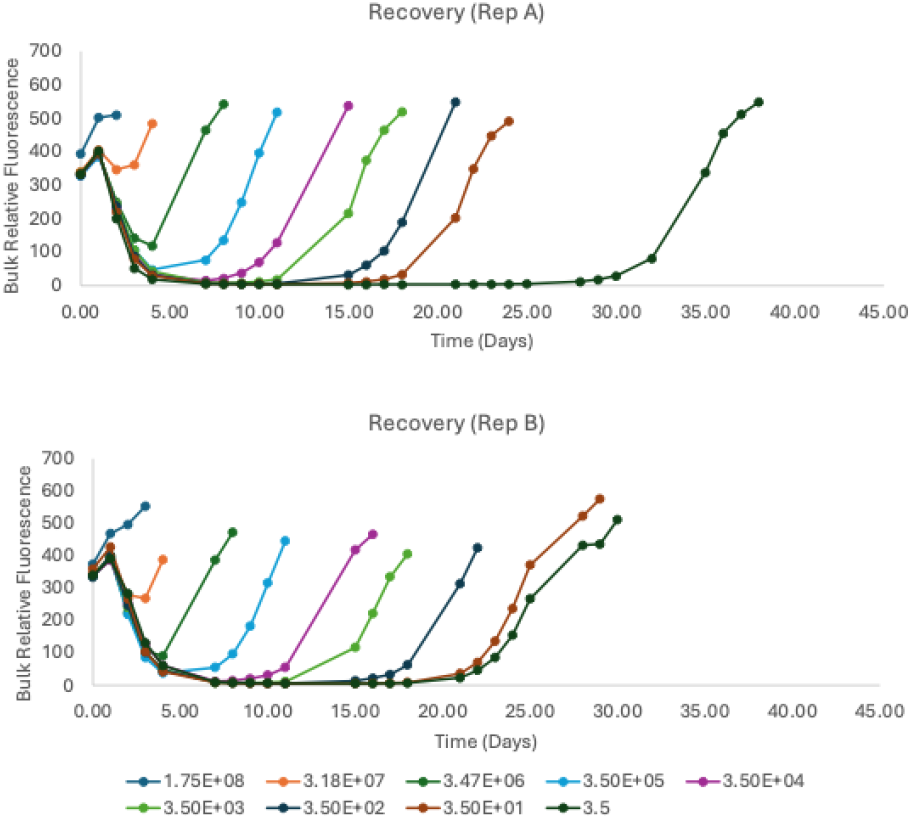
Growth curves for the transformation efficiency and recovery experiments. Colored lines indicate the number of SpecR cells in the inoculum, while the remainder of the 3.5×10^8^ cells/mL is made up of WT cells. Recovery is a function of the SpecR:WT ratio of the inoculum. Two biological replicates are shown.

**Table S1:**
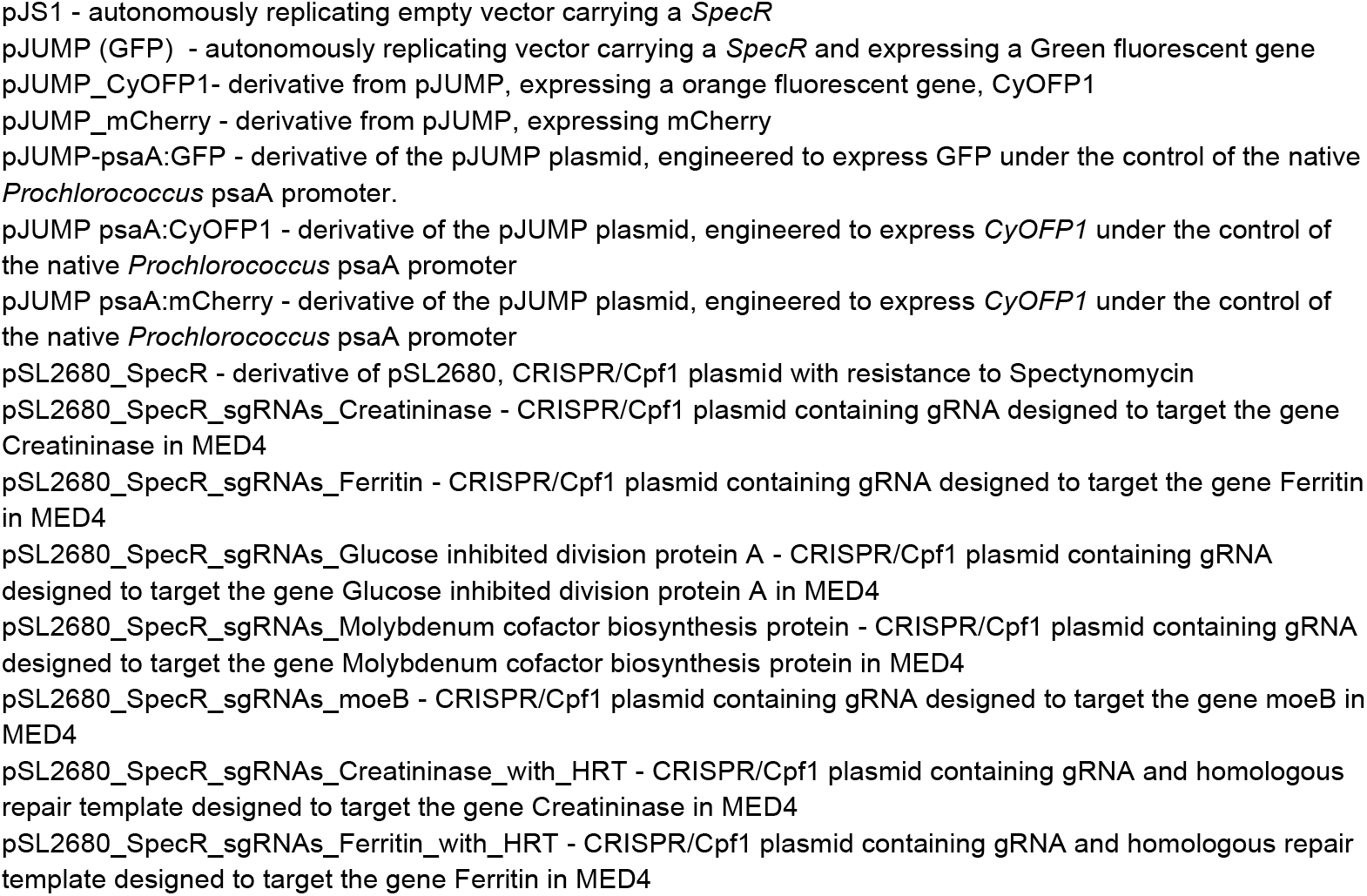
List of plasmids.

**Table S2:**
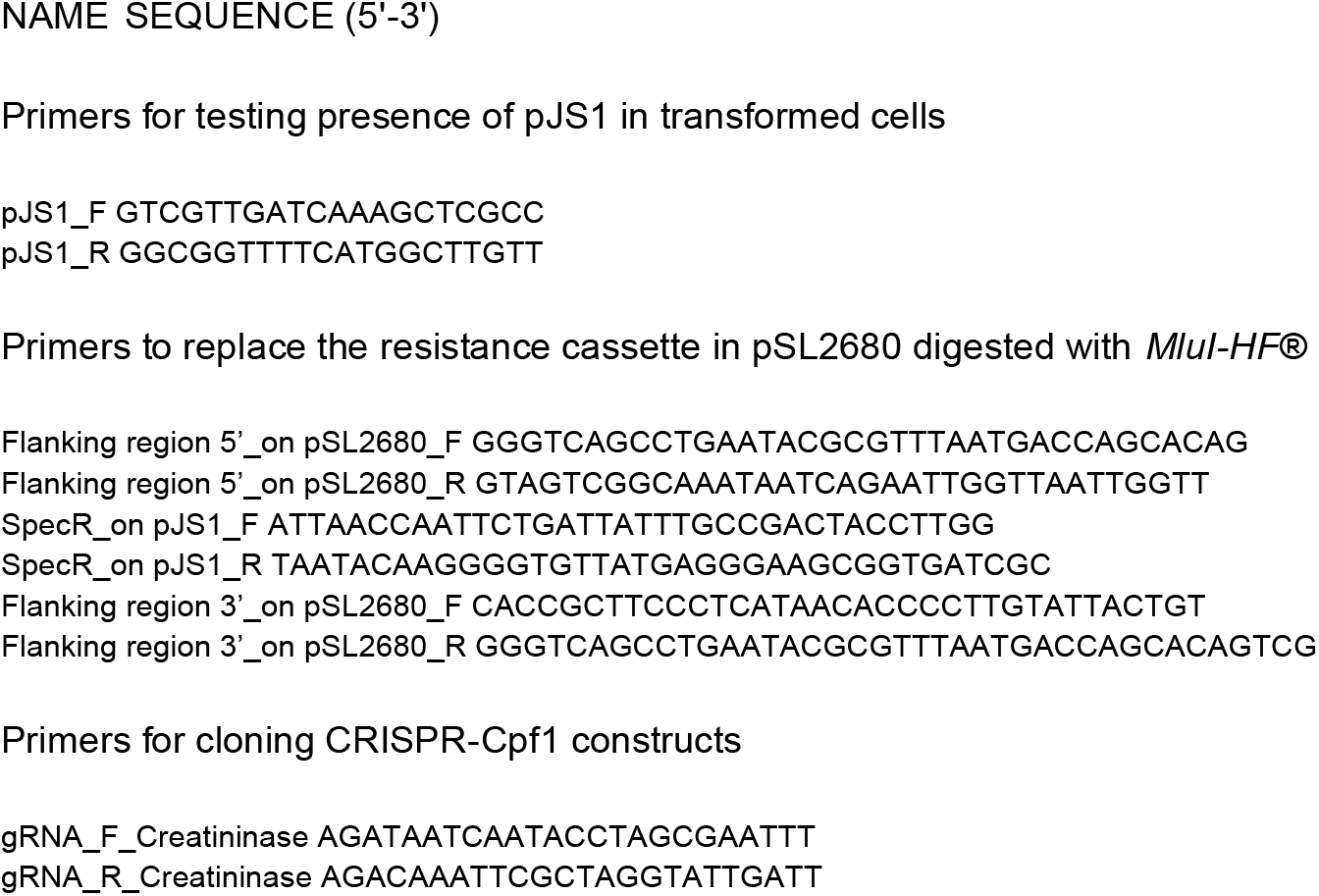
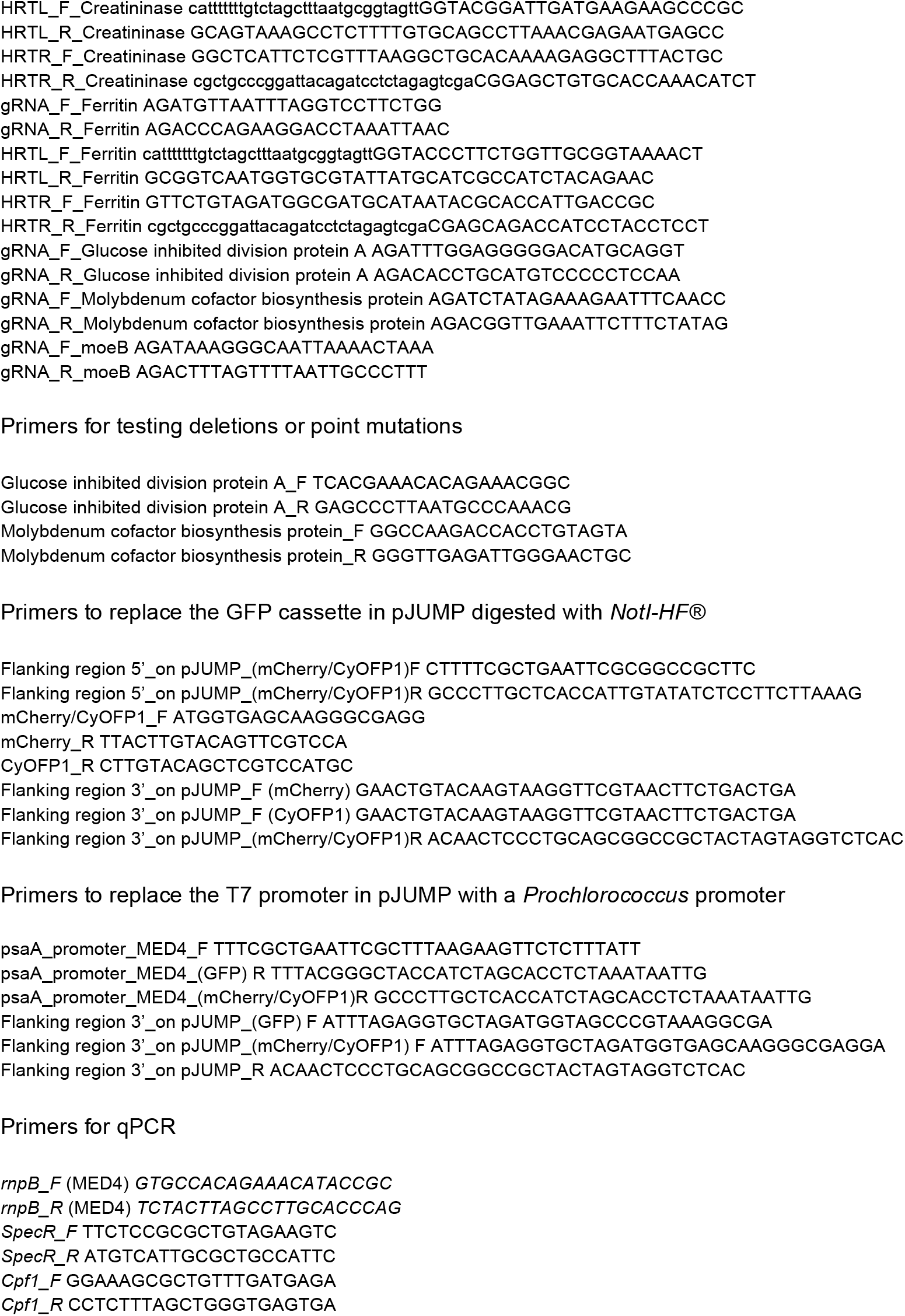
List of primers.

## Notes

### Competing Interest Statement

The authors have declared no competing interest.

